# Helicase Promotes Replication Re-initiation from an RNA Transcript

**DOI:** 10.1101/310235

**Authors:** Bo Sun, Anupam Singh, Shemaila Sultana, James T. Inman, Smita S. Patel, Michelle D. Wang

## Abstract

To ensure accurate DNA replication, a replisome must effectively overcome numerous obstacles on its DNA substrate. After encountering an obstacle, a progressing replisome often aborts DNA synthesis but continues to unwind the DNA, resulting in a gap in the newly replicated DNA. However, little is known about how DNA synthesis is resumed downstream of an obstacle. Here, we examine the consequences of a non-replicating replisome collision with a co-directional RNA polymerase (RNAP). Using single-molecule and ensemble methods, we find that T7 helicase interacts strongly with a non-replicating T7 DNA polymerase (DNAP) at a replication fork. As the helicase advances the fork, the DNAP also moves forward processively, via its association with the helicase. The presence of the DNAP, in turn, increases both helicase’s processivity and unwinding rate. We show that such a DNAP, together with its helicase, is indeed able to actively disrupt a stalled transcription elongation complex, and then initiates replication using the RNA transcript as a primer. These observations exhibit T7 helicase’s novel role in replication re-initiation, independent of replication restart proteins or primase.

## Introduction

Replication fork arrest or collapse occurs when the replisome encounters obstacles such as DNA damage, DNA-bound complexes, or stable DNA secondary structures^1–4^. To avoid replication failure, several mechanisms have evolved to overcome various types of substantial barriers and to restart arrested replication forks independent of replication origins and the existence of these multiple pathways highlights the importance of replication fork recovery^5–8^. However, the mechanisms of fork restart are still unclear. Upon encountering a lesion on the leading strand and in the absence of lesion bypass, a replisome is often found to terminate replication at the lesion, but continues DNA unwinding^9–12^, and this necessitates restarting DNA synthesis downstream from the lesion. A prerequisite for a replisome to resume replication after the lesion is acquisition of a primer, which allows DNA polymerase (DNAP) to re-initiate DNA synthesis. Although re-priming by primase has been shown to rescue replication^13^, this occurs at a relatively low efficiency in overcoming leading-strand lesions and is not adopted by many organisms. There are likely additional replisome recovery pathways to facilitate primer acquisition for replication re-initiation. Replication forks have been shown to pick up primers from stable R-loops or stalled transcriptional elongation complexes^14, 15^. However, the mechanism of this pathway of replication fork restart is not understood.

Here, we investigate this potential replication re-initiation pathway using the bacteriophage T7 replisome. The bacteriophage T7 replisome is a simple model system which provides a powerful *in vitro* system to decipher detailed mechanisms of DNA replication^16–20^. Bacteriophage T7 itself lacks translesion polymerases to perform translesion synthesis directly on lesion sites^21^. However, we recently demonstrated that T7 DNAP, working in conjunction with helicase through specific helicase–DNAP interactions, is able to directly replicate through a leading-strand cyclobutane pyrimidine dimer (CPD) lesion^12^ and this has been observed in other systems^22^. Such a direct lesion bypass occurs in only about 28% of the T7 replisomes, while the remaining population exhibits continued helicase unwinding without DNA synthesis beyond the lesion. This suggests the possible existence of other mechanisms for replication re-initiation downstream of the damage.

In this report, we addressed the questions of whether and how a non-replicating T7 DNAP in the presence of helicase could use a nascent RNA transcript from an RNAP polymerase (RNAP) as a primer to re-initiate DNA replication. We found that a non-replicating T7 DNAP interacts strongly with a T7 helicase at a fork, and this interaction significantly reduces helicase slippage frequency, leading to a faster and more processive unwinding. Furthermore, T7 helicase in association with the non-replicating DNAP is able to displace an RNAP and the DNAP can subsequently re-initiate DNA synthesis using the RNA transcript. These findings reveal a novel pathway for replication re-initiation that is enabled by the participation of helicase.

## Results

### A non-replicating DNAP prevents helicase slippage and increases unwinding rate

During leading-strand replication, DNA synthesis by an actively elongating DNAP has been shown to facilitate T7 helicase unwinding^18, 23, 24^. However, it is unclear if the non-replicating DNAP, which is disengaged from DNA synthesis as could occur after a replisome encountering a lesion, still affects helicase unwinding. Therefore, we first addressed whether a non-replicating DNAP could still facilitate helicase unwinding. Previously, we discovered that T7 helicase, when powered by ATP alone, unwinds DNA rapidly but slips frequently, which is in contrast to its processive unwinding in the presence of dTTP^25^. During a slippage event, helicase loses its grip on the tracking ssDNA, slides backwards under the influence of the re-annealing DNA fork, and then regains its grip to resume unwinding (Fig. 1a). We thus examined T7 helicase’s slippage behavior and unwinding activities (rate and processivity) in the presence of non-replicating DNAP and 2 mM ATP, which is a condition that allows helicase unwinding but does not permit replication. We employed a previously developed single-molecule optical trapping assay to measure T7 helicase unwinding of dsDNA26. Briefly, to mimic a stalled fork with a leading-strand gap, we generated a ssDNA region of approximately 900 nt in the leading strand near a fork junction by mechanically unzipping the dsDNA. Subsequent helicase unwinding of the junction led to an increase in the ssDNA length, allowing tracking of the helicase location (Supplementary Fig. 1a). Once helicase activity was detected, then helicase catalyzed unwinding was monitored under a constant force, which was not sufficient to mechanically unwind the fork junction without a helicase (Fig. 1b). In the absence of DNAP and using only ATP as the fuel source, T7 helicase was found to frequently slip during dsDNA unwinding (Fig. 1b), consistent with our previous findings25. These slippage events led to a remarkable sawtooth pattern in an unwinding trace. The processivity, which is defined as the average distance between slips, was 400 ± 50 bp (mean ± s.d.) under 8 pN of force (Fig. 1d). Surprisingly, we found that in the presence of the non-replicating DNAP (1:1 complex of T7 gp5 and *E. coli* thioredoxin, trx), T7 helicase unwound DNA processively without detectable slippage through the entire dsDNA available (~ 2,500 bp) (Fig. 1c and 1d). Because previous studies showed that T7 DNAP’s functional activities, such as processive synthesis and strong binding on the template, require the association of DNAP with the processivity factor trx27, we investigated whether trx is required to prevent helicase slippage. We found that T7 helicase slipped in the presence of DNAP alone or trx alone but not when both were present, indicating that both DNAP and trx are necessary to prevent helicase slippage (Fig. 1d and Supplementary Fig. 2). For all subsequent experiments with DNAP, trx was also present.

**Figure 1.**
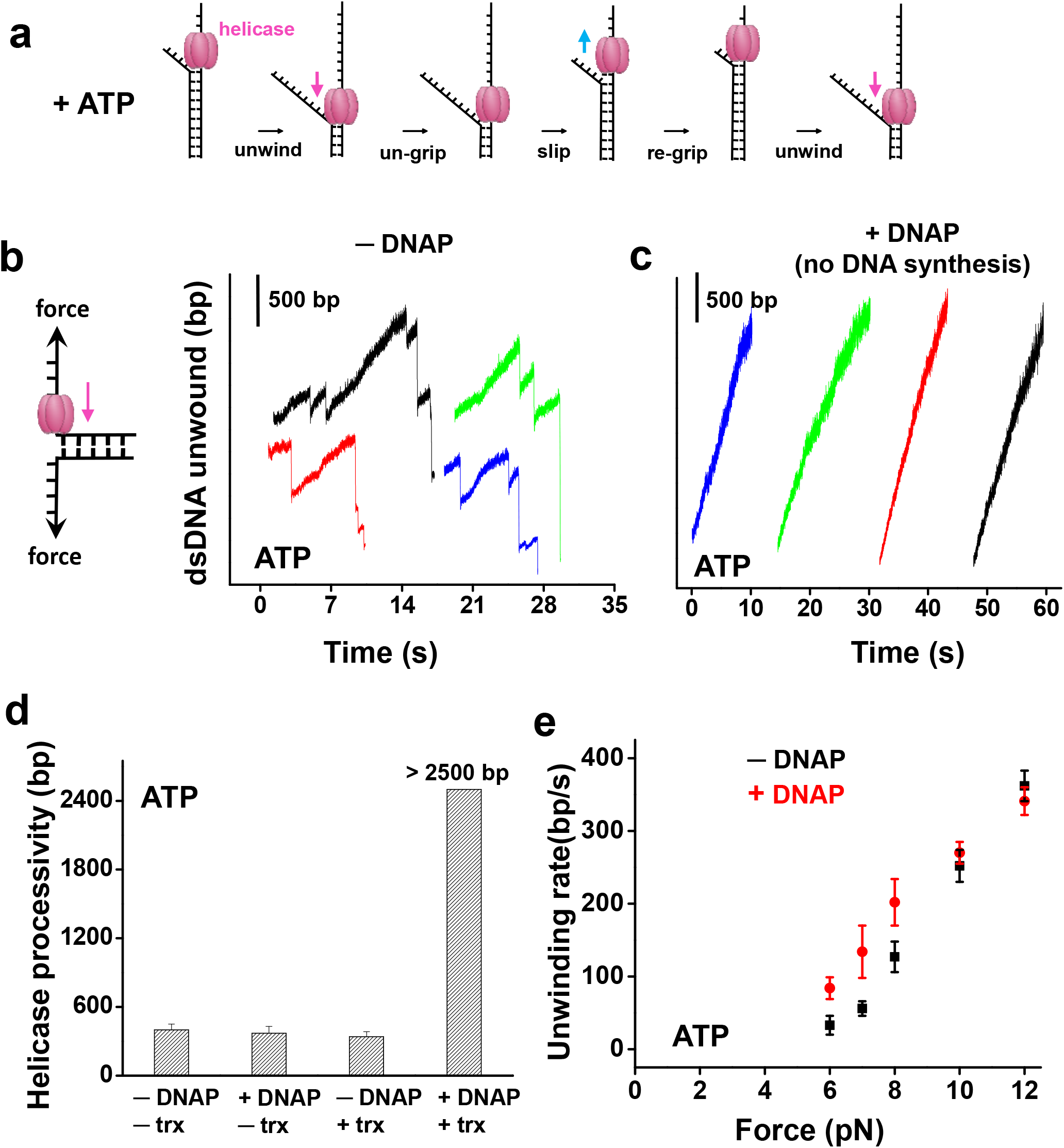
Non-replicating DNAP prevents helicase slippage and increases unwinding rates. **(a)** Cartoon illustrating T7 helicase unwinding and slippage behavior. The helicase unwinds, loses grip, slips, re-grips, and resumes unwinding. Arrows indicate the directions of helicase movements. **(b)** and **(c)** Representative traces showing the number of unwound base pairs by T7 helicase versus time in the absence or presence of non-replicating DNAP, respectively. Experiments were conducted with 2 mM ATP under 8 pN force. For clarity, traces have been arbitrarily shifted along both axes. **(d)** Measured processivity of T7 helicase (mean distance between slippage events) in the absence and presence of DNAP and/or trx with 2 mM ATP under 8 pN external force. **(e)** T7 helicase unwinding rates as a function of force in the absence or presence of the DNAP with 2 mM ATP. Unwinding rates were determined between slippage events (if any).

Next, we examined how the stimulation of helicase unwinding by a non-replicating DNAP depended on the force. At forces below 9 pN, helicase alone unwound with frequent slippage, and DNAP increased the helicase unwinding rates between slips, whereas this effect was negligible in a higher force region (10 to 12 pN) (Fig. 1e). A similar trend in unwinding rates was observed in experiments carried out in the presence of dTTP (Supplementary Fig. 3). Therefore, we anticipate that in the absence of force, helicase’s unwinding activities may also be stimulated by the presence of a non-replicating DNAP.

To examine whether the regulation of helicase unwinding activities by a non-replicating DNAP is via their direct interactions, we utilized a mutant T7 helicase that lacks the 17 carboxyl-terminal amino acid residues (ΔCt) required for interaction with T7 DNAP^28 29^. The ΔCt mutant of T7 helicase has previously been shown to have helicase unwinding capability, but does not form a stable complex with DNAP^28^. We carried out similar unwinding experiments to those described above except with the ΔCt mutant. We found that slippage occurred both in the presence and absence of a non-replicating DNAP. The non-replicating DNAP did not change the processivity or the unwinding rate of the mutant helicase (Supplementary Fig. 4), which is in stark contrast to the results observed with wt helicase (Fig. 1). Therefore, direct interactions between a helicase and a non-replicating DNAP are essential for the observed slippage prevention and unwinding rate enhancement.

### A non-replicating DNAP with a helicase localizes to the fork junction

The finding that a non-replicating DNAP facilitates helicase unwinding raises question about the configuration of the helicase and the DNAP at the fork. If DNAP is able to bind to the leading strand while interacting with helicase on the lagging strand across the fork junction, then it will be poised for leading-strand synthesis once it acquires a primer (Fig. 2a). We therefore designed an experiment to investigate whether a non-replicating DNAP binds to the leading strand at the fork junction during helicase unwinding. First, in the presence of helicase, dTTP, and with or without DNAP, we mechanically unzipped approximately 900 bp of a 4,100 bp dsDNA template (at a speed of 250 bp/s) (step 1). This created a ssDNA region for helicase loading and allowed subsequent translocation of the helicase to the fork. Helicase was then monitored as it unwound under a constant force of 9 pN (step 2). Finally, once helicase unwound approximately 1000 bp, we rapidly unzipped the remaining DNA mechanically at 2,000 bp/s (much faster than helicase unwinding rate) to detect any bound proteins across the fork junction (step 3). During this mechanical unzipping step, a force rise significantly above the naked DNA baseline served as a sensitive detector for the presence of a bound protein across of the fork junction^30–33^.

**Figure 2.**
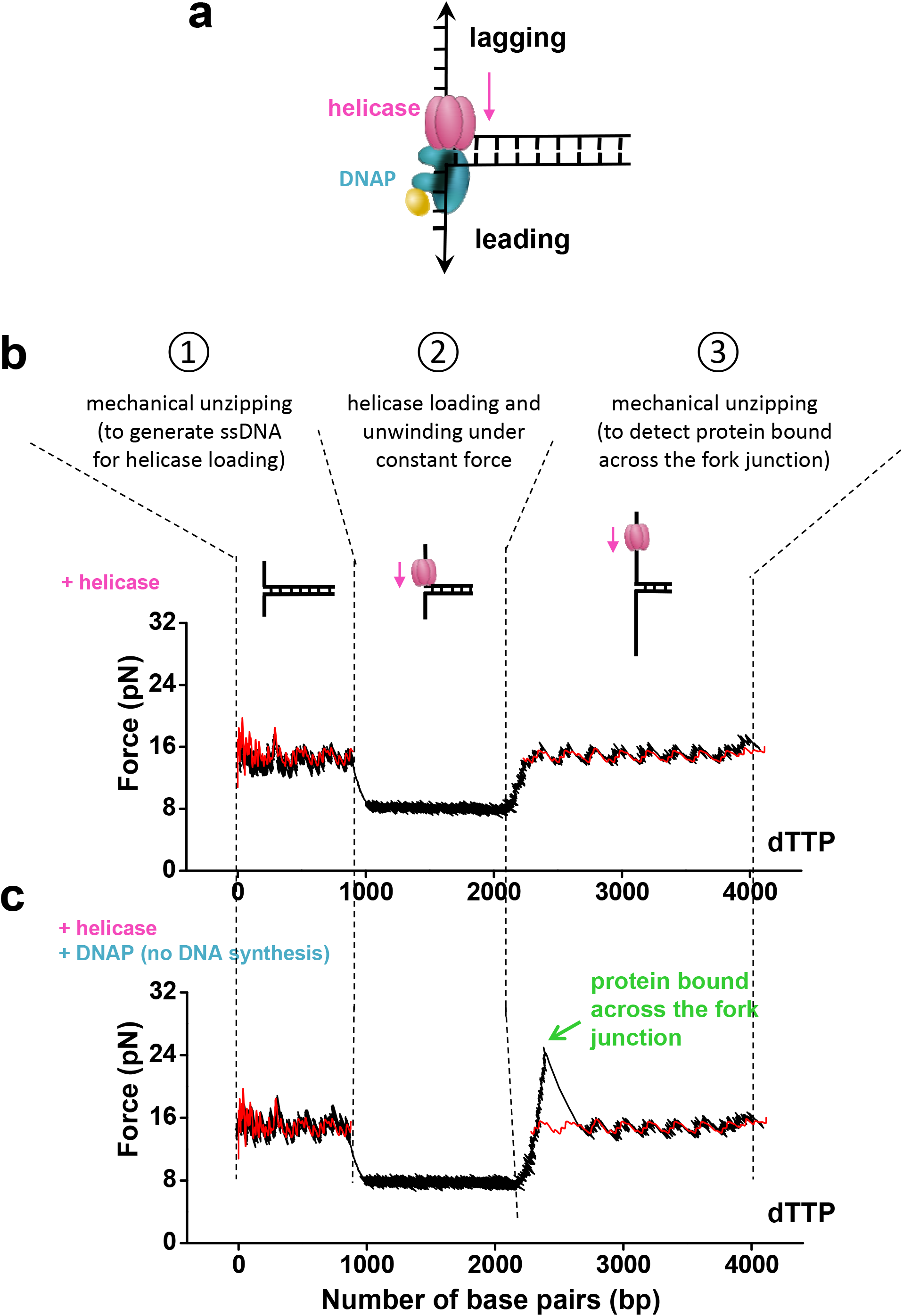
Non-replicating DNAP localizes to the fork with the helicase. **(a)** A cartoon illustrating helicase-DNAP coupling. A non-replicating DNAP directly interacts with an unwinding helicase. The helicase-DNAP complex is juxtaposed crossed the fork junction. **(b)** and **(c)** Representative traces showing the force versus number of base pairs unzipped/unwound in presence of helicase and 2 mM dTTP, either without or with the presence of non-replicating DNAP respectively. The red curves correspond to unzipping naked DNA. The green arrow indicates a force peak above the naked DNA baseline.

In the absence of DNAP, no significant force rise above the naked DNA baseline was detected during step 3 (Fig. 2b), suggesting that helicase translocates on the lagging-strand and unwinds DNA while having minimal interactions with the leading strand. In contrast, in the presence of non-replicating DNAP, in step 3, about 75% of the traces exhibited a force rise significantly above the naked DNA baseline, followed by a return of the unzipping force to the naked DNA baseline (Fig. 2c). This demonstrates that a protein or protein complex was located across the fork junction in those traces. The peak force was 26 ± 1 pN (mean ± s.d.) (Supplementary Fig. 5), whereas the naked DNA baseline averaged to approximately 15 pN. The absence of a force peak under the helicase only condition (Fig. 2a) precludes the possibility that this force peak is due to interactions of helicase alone with the DNA fork junction. The force peak was also absent in a control experiment with DNAP and ΔCt helicase (Supplementary Fig. 6), suggesting that direct interactions between DNAP and helicase were essential. The force rise is thus attributed to the helicase/DNAP interactions across the fork junction. For the 25% of traces that did not show detectable force rise above the naked DNA baseline, it is possible that DNAP was not present at the fork junction. However, because we observed a complete lack of slippage during helicase unwinding in the presence of DNAP (Fig. 1), we favor the possibility that DNAP was present at the fork but the DNAP-helicase interactions were transiently lost at the moment of detection. Ultimately, we conclude that a non-replicating DNAP directly interacts with an unwinding helicase right across the fork junction (Fig. 2a).

### Helicase assists a non-replicating DNAP in displacing RNAP and re-initiating replication

A non-replicating DNAP localized at a fork junction via an unwinding helicase is well poised for re-initiating replication upon primer acquisition. This may take place during a co-directional collision with a transcription elongation complex (TEC) if a helicase-DNAP complex is able to displace the RNAP and allow DNAP access to the RNA transcript^7^. Although we have demonstrated that a non-replicating DNAP enhances helicase’s motor ability (Fig. 1), it is unclear whether the non-replicating helicase-DNAP complex is capable of overcoming the TEC barrier.

To investigate the outcome of such a collision, we developed a real-time assay to monitor co-directional progression of a helicase with a non-replicating DNAP through a stalled *E. coli* TEC (Supplementary Fig. 1b). During an experiment, the reaction buffer contained non-replicating DNAP and all four dNTPs to permit potential transition to DNA synthesis, while the extension was monitored under a low constant unzipping force. To differentiate between unwinding without and with DNA replication, we first carried out two control experiments to characterize the rate of extension. In the first experiment, the leading strand was provided with a DNA primer containing a 3’ inverted dT from which T7 DNAP could not synthesize (Supplementary Fig. 1b). We observed that the DNA length increased at a rate of 34 ± 20 nm/s (mean ± s.d) (Fig. 3a). In the second experiment, when the leading strand was provided with a DNA primer from which T7 DNAP could synthesize, the DNA length increased at a rate of 91 ± 18 nm/s (Fig. 3a). Thus, a transition to this faster rate serves as a clear indicator for onset of DNA synthesis.

**Figure 3.**
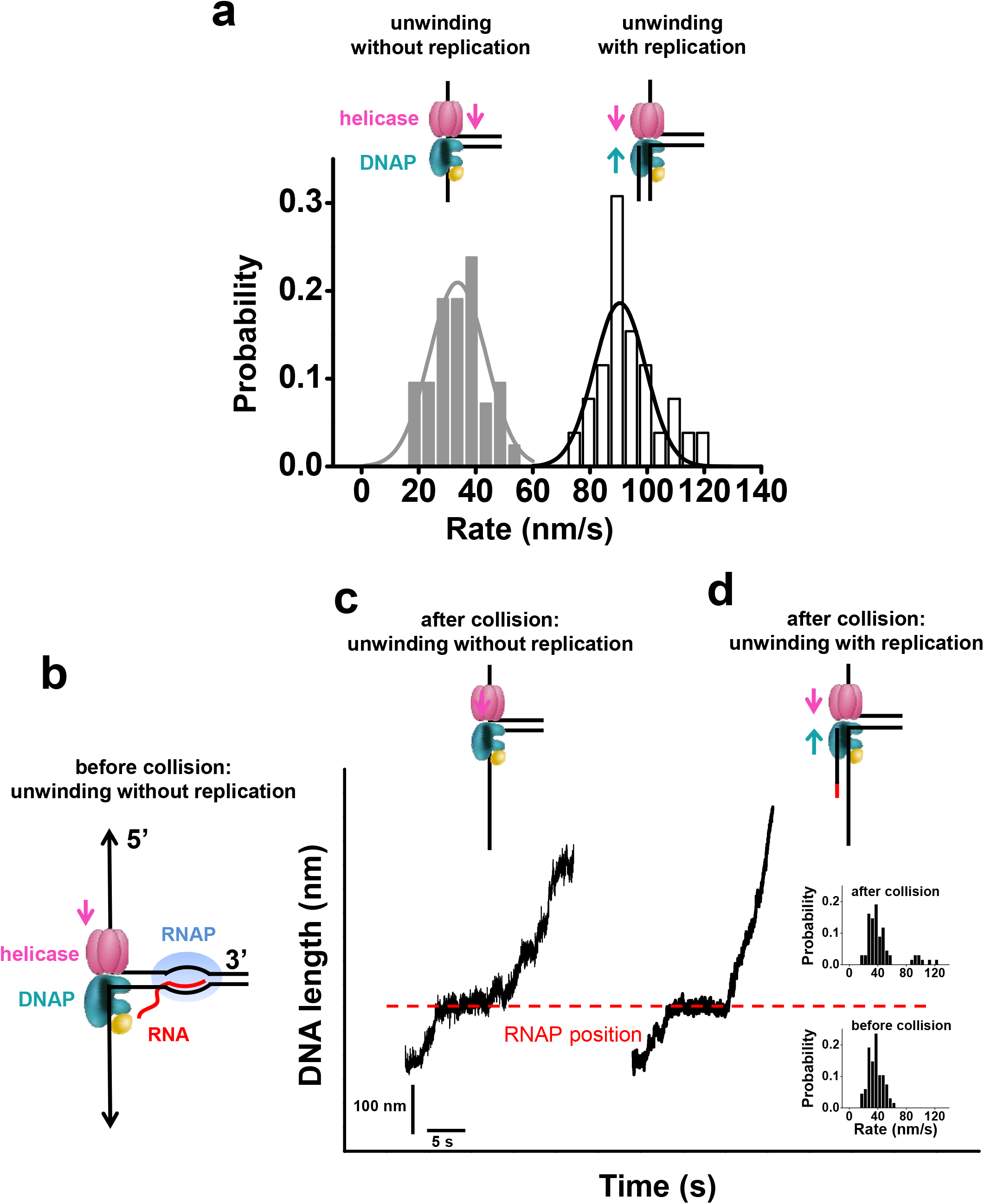
Single molecule experiments on helicase and non-replicating DNAP collision with a TEC. **(a)** Distributions of DNA length increase rates of helicase unwinding with or without replication in 0.5 mM dNTP (each) under 5 pN of force. **(b)** Experimental design. Left panel shows a cartoon of *E. coli* TEC that was stalled at + 20 nt from the promoter while a helicase with a non-replicating DNAP encountered the TEC co-directionally. **(c)** A representative trace of unwinding without replication after collision. Experiments were carried out in the presence of helicase, DNAP, and 0.5 mM dNTP (each) under 5 pN of force. The dotted line indicates the expected stalled TEC position. The cartoon on the top illustrate replication status after the collision with RNAP. **(d)** A representative trace of unwinding with replication after collision. Same experimental conditions were used as in (c). For clarity, traces have been shifted along the time axis. Insets display the distributions of DNA length increase rates before and after the collision.

To directly examine the consequences of collision between the helicase/non-replicating DNAP and a TEC, we used a parental DNA template which contained no pre-existing DNA primers but a co-directional TEC stalled at +20 nt from its promoter (Fig. 3b). Thus, there should be only unwinding without DNA synthesis prior to their collision. Consistent with this, all traces exhibited an expected slow DNA length increase rate before collision (Fig. 3c and 3d). Upon collision, several types of behavior were observed. In 88% of 68 traces, DNA length continued to increase at a rate similar to that before collision (Fig. 3c), consistent with unwinding without DNA synthesis. Within this 88%, 43% of the traces showed a detectable pause at the expected RNAP position (Fig. 3c), suggesting that the helicase-DNAP complex was able to overcome the TEC roadblock but the complex moved forward without DNA synthesis; while the rest did not show any detectable pausing, in part due to an absence of a TEC during the initial TEC formation (Methods). Interestingly, the remaining 12% of traces paused transiently at the expected RNAP position and then transitioned to an increased rate consistent with that of unwinding with leading-strand synthesis (Fig. 3d). For these traces, the helicase-DNAP complex was indeed able to overcome the TEC roadblock, and the DNAP was then able to carry out DNA synthesis. Normalizing against the initial TEC formation efficiency, the replication re-initiation efficiency is approximately 15%.

These results demonstrate synergistic coordination between an unwinding helicase and a non-replicating DNAP. While a non-replicating DNAP greatly enhances helicase unwinding activities, helicase interactions in turn keep a DNAP close to the unwinding fork and promote the DNAP in re-initiating leading-strand synthesis. We conclude that helicase in association with a non-replicating DNAP forms a strong motor complex at the fork, capable of displacing an RNAP. As a result, DNAP can gain access to the RNA transcript and use it as a primer for replication initiation.

### Ensemble studies support that helicase promotes replication re-initiation

To exclude the possibility that the observed replication re-initiation was a result of the force in the ssDNA under our single molecule conditions, we carried out corresponding ensemble experiments in the absence of any externally applied force. For these experiments, the replication fork substrate contained a stalled, co-directional T7 TEC on the parental dsDNA (Fig. 4a and Supplementary Fig. 7). The TEC was able to make run-off products with rNTPs, (Supplementary Fig. 7), and indeed T7 DNAP alone was able to efficiently extend the RNA primer on ssDNA template and make run-off products (Fig. 4c). However, no run-off products were observed with dNTPs on a fork substrate when helicase was absent (Fig. 4a). The TEC was able to use dNTPs^34^, but was only able to incorporate a few nucleotides and these short dNTP-elongated RNA primers migrated on the gel close to the 10-mer DNA markers (Fig. 4a and Supplementary Fig. 8). We observed run-off products in the presence of helicase, DNAP, and dNTPs and we estimated that ~ 14% of the RNA had fully extended to the end within 10 minutes (Figs. 4a and 4d). These experiments in combination indicate that the run-off products result from replication re-initiation by T7 DNAP using the RNA as a primer, and helicase plays an essential role in replication re-initiation on the fork substrate. As additional control experiments, we replaced the fork substrate with a blunt end dsDNA, which T7 helicase cannot load onto and is thus unable to unwind (Fig. 4b)^35^. As with the forked substrate, the blunt dsDNA template contained a stalled TEC (Supplementary Fig. 7). In the presence of dNTPs only, the TEC was only able to transcribe a few nucleotides. When helicase and DNAP were also present, fully extended primers were not detected on this blunt substrate (Fig. 4b and 4d), in contrast to the fork substrate. Thus helicase unwinding is critical in enabling re-initiation.

**Figure 4.**
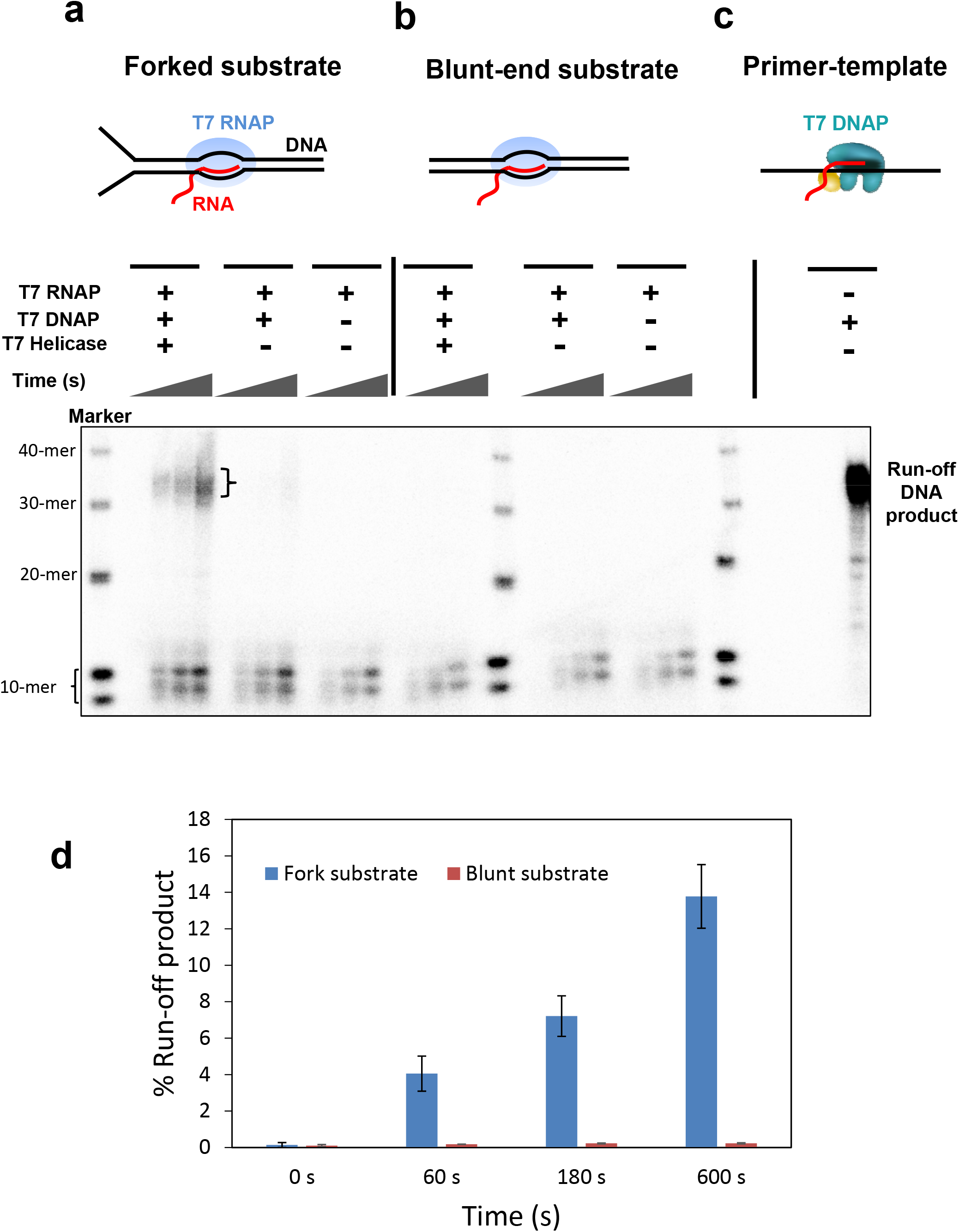
Bulk experiments on helicase and non-replicating DNAP collision with a TEC. **(a)** and **(b)** Experiments on fork DNA substrate (left) and blunt DNA substrate (right) respectively. The parental DNA contained a stalled T7 RNAP and experiments were carried out with dNTP mixture spiked with [α-^32^P]-dGTP. For each experiment, reactions were quenched at four time points (0, 60, 180 and 600 s). Samples were mixed with formamide and bromophenol blue dye and heated at 95°C for five minutes before loading on 12% acrylamide/6 M urea sequencing gels. Sequencing gels show the kinetics of the RNA primer extension on either the fork substrate or the blunt substrate. The run-off DNA product is 38-nt long. The products running close to the 10-nt DNA markers are 14-mer and 15-mer resulting from dNTPs addition by T7 RNAP to the 12-mer RNA primer. **(c)** Replication reaction performed with just the primer annealed to the template. Experiment was carried out with dNTP mixture spiked with [α-^32^P]-dGTP and T7 DNAP. Reaction was quenched at 600 s. The replication product was used as a control for quantitating the % run-off DNA products obtained in (a) and (b). **(d)** % run-off product estimated from (a) and (b).

These results are in agreement with our single-molecule findings that a non-replicating DNAP, in conjunction with an unwinding helicase, can utilize an RNA primer from a co-directional TEC and subsequently carry out continuous DNA synthesis. These data also reinforce the conclusion that helicase is essential in assisting the DNAP in re-initiating the leading-strand synthesis.

## Discussion

Although the long established role of replicative helicases is to catalyze strand separation, emerging evidence now supports the notion that their functions in replication are much broader^36, 37^. Our findings here elucidate a novel role of T7 helicase in enabling replication re-initiation (Fig. 5). Recent structural studies have shown interactions between the T7 helicase and T7 DNAP^29, 38^. Here we show that during DNA unwinding, helicase strongly interacts with a non-replicating DNAP on the leading strand. The two proteins form a complex to clamp across the fork junction and their interactions permit the DNAP to travel alongside the unwinding helicase. The presence of the DNAP at the fork increases helicase’s unwinding rate and processivity by preventing helicase slippage. Consequently, processive unwinding of the helicase in association with DNAP leads to TEC disruption during a co-directional collision with transcription, exposing the RNA transcript. Then the DNAP is able to pick up the RNA and use it as a primer to initiate DNA synthesis. Although previous work showed that a *replicating* replisome involving a T4 or *E. coli*. leading-strand DNAP may overcome a TEC barrier^14, 39, 40^, this work shows that even if the T7 DNAP is not replicating, T7 helicase can enhance its capacity to overcoming barriers and re-initiating replication.

**Figure 5.**
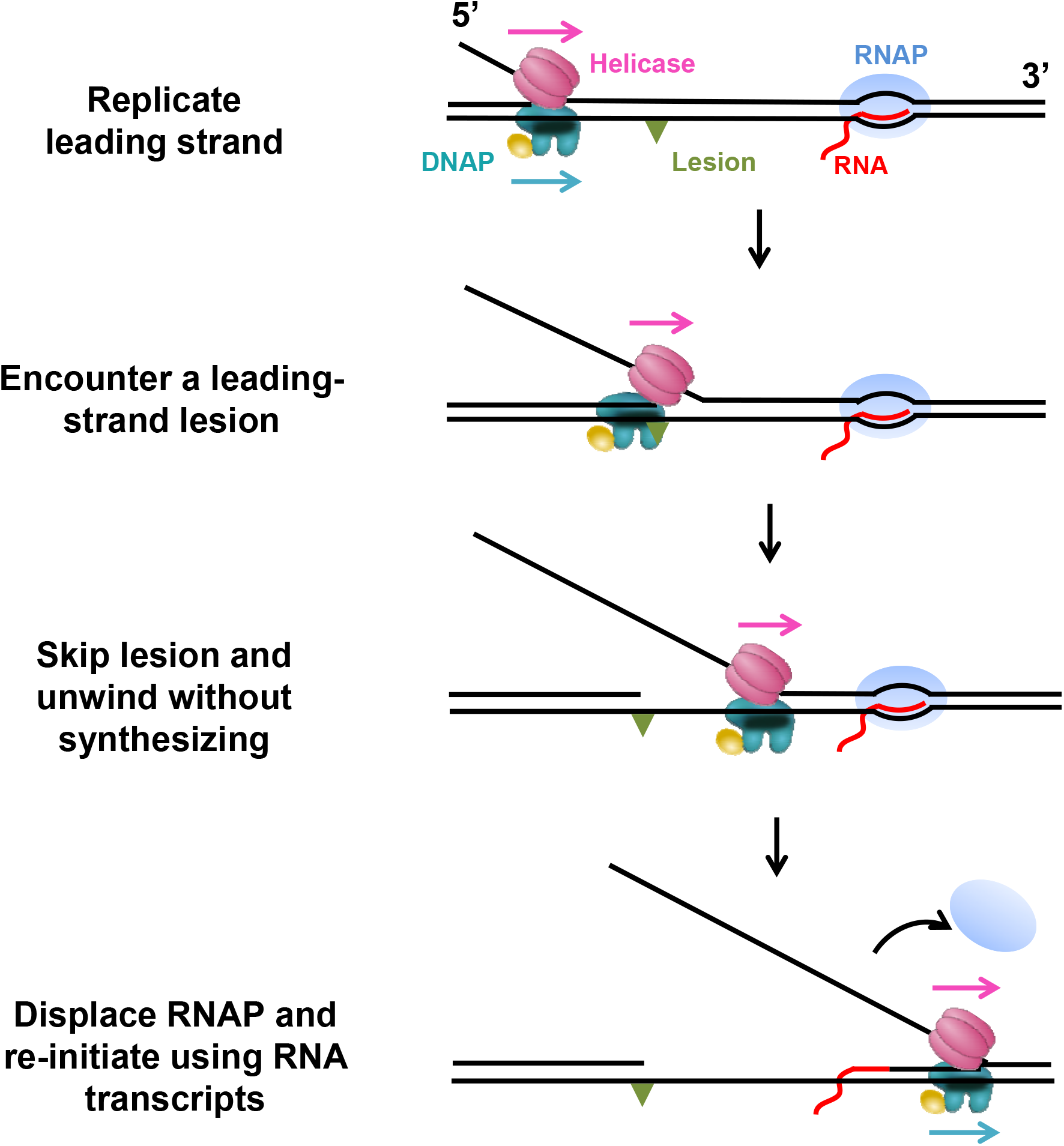
Proposed T7 replication re-initiation model. Cartoons illustrate a proposed model for T7 replication re-initiation. The replisome here consists of leading-strand DNAP and helicase. When the replisome encounters a leading-strand lesion, helicase may continue to unwind processively via association with a non-replicating DNAP. The two proteins form a complex clamping crossed the fork junction, poising the DNAP for replication re-initiation. After collision with a TEC, the helicase-DNAP displaces the RNAP and the DNAP then uses the RNA as a primer to re-initiate the replication.

This work may also provide insight to transcription-initiated replication found in T7, T4, *E. coli* ColE1 plasmid, mitochondrial DNA replication, and origin-independent replication initiation in eukaryotes, whose mechanisms remain elusive41. In T7, it was found that helicase enhances DNAP’s acquisition of RNA, from T7 RNAP, to use as a primer42-45. In this process, DNAP must be positioned in close vicinity to the T7 RNAP; however, the DNAP, without DNA synthesis, has low affinity to dsDNA. Thus, we can envision that DNAP’s association with helicase may be highly beneficial to initiation by bringing the DNAP to the origin of the replication, forming a strong complex to displace the RNAP, and picking up a primer from R-loops.

Our work supports that T7 helicase not only participates in the assembly of the replication machinery at the fork, but also assists in replication conflict resolution with roadblocks. Thus helicase has broad functionalities and unexpected roles in replication.

## Methods

### Protein and DNA preparations

Untagged gp4A’ (wild-type helicase) and delta C gp4A’ mutant (ΔCt) were overexpressed in the BL21 DE3 cell line^28, 46^. Untagged T7 gp5 (D5A, D7A) was overexpressed in the A179 cell line containing pGP1 plasmid47. The cells were lysed by three freeze thaw cycles in 20 mM phosphate buffer pH 7.4, with 50 mM NaCl, 10% glycerol, 2 mM DTT, 2 mM beta mercapto-ethanol and 1 mM or 2.5 mM EDTA (for the helicase or the gp5 respectively) in the presence of 0.2 mg/ml lysozyme. Polyethyleneimine precipitation was carried out by increasing the salt concentration to 0.5 M. The supernatant was precipitated in 70 % ammonium sulfate, and purified by Phosphocellulose (P11 resin) followed by DEAE Sepharose column chromatography. *E. coli* thioredoxin (trx) was purchased from Sigma-Aldrich (St. Louis, MO). *E. coli* RNAP was expressed and purified as previously described at a low level in wild type *E. coli* strains (WT, DH5α) to yield RNAPs^48^. *E. coli* RNAPs were purified to homogeneity by using a modification of the method of Burgess and Jendrisak to include chromatography on nickel agarose^49^. T7 RNAP was overexpressed in *E. coli* strain BL21/pAR1219 and purified using three chromatographic columns consisting of SP-Sephadex, CM-Sephadex, and DEAD- Sephacel^50, 51^. The purified enzyme was dialyzed against buffer (20 mM sodium phosphate, pH 7.7, 1 mM Na3-EDTA, and 1 Mm DTT) containing 100 mM NaCl and 50% (v/v) glycerol, and stored at -70 °C.

The DNA template for helicase unwinding and unzipping consisted of a 1.1 kbp anchoring segment and a 4.1 kbp unwinding segment (Supplementary Fig. 1a)^52^. In brief, the anchoring segment was amplified from plasmid pRL574^53^ and the unwinding segment was derived from 17 pseudo-repeats (or 17mer) of the 5s rRNA sequence^54^. The final product was produced by ligating the two segments in a 1:1 molar ratio. The DNA template for the helicase-DNAP coupled replication initiation assay consisted of three pieces: two arms and a trunk (Supplementary Fig. 1b). Arm 1 (1129 bp) was PCR amplified from plasmid pRL574 using a digoxigenin-labeled primer. The resulting DNA fragment was digested with BstXI (NEB, Ipswich, MA) to create an overhang and was subsequently annealed to a short DNA with a complementary overhang formed by adapter 1 (5’-/phos/GCA GTA CCG AGC TCA TCC AAT TCT ACA TGC CGC-3’) and adapter 2 (5’-/phos/GCC TTG CAC GTG ATT ACG AGA TAT CGA TGA TTG CG GCG GCA TGT AGA ATT GGA TGA GCT CGG TAC TGC ATCG-3’). Arm 2 (2013 bp) was PCR amplified from plasmid pBR322 (NEB, Ipswich, MA) using a biotin-labeled primer. The resulting DNA fragment was digested with BstEII (NEB, Ipswich, MA) to create an overhang and was subsequently annealed to adapter 3 (5’-/phos/GTAAC CTG TAC AGT GTA TAG AAT GAC GTA ACG CGC AAT CAT CGA TAT CTC GTA ATC ACG TGC AAG GC CTA-3’). The adapter 3 from arm 2 and the adapter 2 from arm 1 were partially complementary to each other and were annealed to create a short ~30-bp trunk with a 3-bp overhang for the trunk ligation. The 1.5 kbp trunk containing the T7A1 promoter was amplified from plasmid pRL574. The final product was produced by ligating the arms with the trunk at 1:4 ratio.

### Single-molecule assays

Sample chamber preparation was similar to that previously described^12, 25, 52^. Briefly, DNA tethers were formed by first non-specifically coating a sample chamber surface with anti-digoxigenin (Roche, Indianapolis, IN), which binds nonspecifically to the coverglass surface, followed by an incubation with digoxigenin-tagged DNA. Streptavidin-coated 0.48 mm polystyrene microspheres (Polysciences, Warrington, PA) were then added to the chamber. DNAP was assembled by adding 10 μM of gp5 in 50 μM *E. coli* trx and incubating at room temperature for 5 minutes. The helicase and DNAP were prepared as follows: first, 100 nM of the appropriate helicase hexamer was incubated for 20 min in the replication buffer on ice; then 100 nM of the appropriate DNAP was added and the solution was incubated for 10 min at room temperature. This solution was then further diluted to obtain the final experimental concentrations of helicase and DNAP, nucleotides and MgCl_2_. The resulting solution was flowed into the chamber just prior to data acquisition. The unwinding and replication buffer consisted of 50 mM Tris-HCl (pH 7.5), 40 mM NaCl, 10% glycerol, 1.5 mM EDTA, 2 mM DTT and dNTPs at the concentrations specified in the text, and MgCl_2_ at a concentration of 2.5 mM in excess of the total nucleotide concentration. Paused transcription complexes were formed by incubating 2 nM *E. coli* RNAP, 0.4 nM DNA template and 1 mM ApU, and 1 mM ATP/CTP/GTP in transcription buffer for 30 min at 37°C^31^.

Experiments were conducted in a climate-controlled room at a temperature of 23.3°C, but owing to local laser trap heating the temperature increased slightly to 25 ± 1°C^55^. Each experiment was conducted in the following steps. First, several hundred base pairs of dsDNA were mechanically unzipped, at a constant velocity of 1,400 bp/s (helicase unwinding assay) or 250 bp/s (DNAP binding detection assay), to produce a ssDNA loading region for helicase. Second, DNA length was maintained until a force drop below a preset value, indicating helicase unwinding of the DNA fork. Third, a constant force was maintained at this preset value via computer-controlled feedback, while helicase unwound the dsDNA. In the unzipping force analysis of helicase and DNAP association, after the detection of helicase loading and unwinding, the remaining dsDNA was mechanically unzipped at an extremely fast velocity of 2,000 bp/s to probe the potential interactions at the fork.

### Data collection and analysis

Data were low-pass filtered to 5 kHz and digitized at 12 kHz, then were further averaged to 110 Hz. The acquired data signals were converted into force and DNA extension as previously described26. In the helicase-unwinding studies, one separated base pair generated two nucleotides of ssDNA. Accordingly, real-time DNA extension in nm was further converted into the number of base pairs unwound based on the elastic parameters of ssDNA under our experimental conditions. To improve positional accuracy and precision, the data were then aligned to a theoretical unzipping curve for the mechanically unzipped section of the DNA. Helicase unwinding rates were determined as previously described26. For the helicase-DNAP coupled replication initiation assay, we had to determine whether the movement after the RNAP was due to helicase alone or helicase-coupled DNAP synthesis and this was more readily achieved by directly measuring the DNA length increase rate in nm/s. Therefore, we presented data as DNA length in nm and rates in nm/s. The position of a paused TEC was known from the DNA template design (753 bp from the initial fork.) Its position in nm showed in Fig. 3 was determined by converting bp to nm.

### Ensemble assays

For experiments described in Fig. 4, fork substrate and blunt substrate (sequences provided in Supplementary Fig. 7a) were annealed by mixing template, non-template, and 5’-fluorescein labeled RNA primer in 1.25:1.5:1 ratio in Tris-HCl pH 7.5, 1 mM EDTA, 50 mM NaCl buffer, incubating the mixture at 95 °C for 2 minutes and gradually cooling it to 20 °C. The TEC was assembled by mixing the substrate (200 nM) with T7 RNAP (1100 nM), T7 helicase (220 nM), and dTTP (1 mM) at 18 °C for 60 minutes in buffer containing Tris-HCl pH 7.5, 40 mM NaCl, 10% glycerol, 2 mM EDTA and 2 mM DTT. Assembled TEC was then incubated with T7 DNAP (220 nM) and thioredoxin (1100 nM) for 60 minutes. Reactions were initiated by adding the rest of the dNTPs (200 μM each, spiked with [α-32P]-dGTP) and MgCl2 (final concentration 5 mM in the reaction). After preset time intervals (0, 60, 180 and 600 s), the reactions were stopped with EDTA (0.15 M), mixed with formamide and bromophenol blue dye, heated at 95°C for five minutes and loaded on a 12% polyacrylamide/6 M urea sequencing gels. Gels were exposed to phosphor screens and the screens were scanned with Typhoon FLA 9500 scanner (GE Healthcare). Replication reaction was also performed with just the primer annealed to the template. The replication product of this reaction at 600 s was used as a control for quantitating the percent run-off DNA products.

## Acknowledgements

We thank Dr. Robert M. Fulbright Jr. for purification of the *E. coli* RNA polymerase, Dr. Manjula Pandey for purification of T7 helicase and T7 DNAP, and Dr. Shanna Moore and the Sun laboratory for critical reading of the manuscript. We wish to acknowledge support from, National Key Research and Development Program of China (2016YFA0500900 and 2017YFA0106700 to B.S.), Shanghai Pujiang Program (16PJ1406900 to B.S.), National Institutes of Health grants (GM059849 to M.D.W. and R35GM118086 to S.S.P.), Howard Hughes Medical Institute, and National Science Foundation grant (MCB-1517764 to M.D.W.).

## Author Contributions

B.S. and M.D.W. designed the single molecule experiments. B.S. carried out the single-molecule experiments, and analyzed and interpreted single-molecule data. J.T.I. maintained and upgraded the optical trapping setup. A.S., S.S., and S.S.P. designed the ensemble replication assays. S.S. and A.S. performed ensemble experiments. B.S. and M.D.W. drafted the manuscript. All authors contributed to revisions of the manuscript and intellectual discussions.

